# Increasing adult density compromises anti-bacterial defense in *Drosophila melanogaster*

**DOI:** 10.1101/2022.01.02.474745

**Authors:** Paresh Nath Das, Aabeer Basu, Nagaraj Guru Prasad

**Author notes:** Author contributions: Conceptualization & Methodology, PND and AB; Investigation & Data curation, PND and AB; Formal analysis, Visualization & Writing (first draft), PND and AB; Writing (review and editing), PND, AB, and NGP; Funding acquisition, NGP. Declaration of interests: The authors declare no competing interests.

## Abstract

The density-dependent prophylaxis hypothesis predicts that risk of pathogen transmission increases with increase in population density, and in response to this, organisms mount a prophylactic immune response when exposed to high density. This prophylactic response is expected to help organisms improve their chances of survival when exposed to pathogens. Alternatively, organisms living at high densities can exhibit compromised defense against pathogens due to lack of resources and density associated physiological stress; the density stress hypothesis. We housed adult *Drosophila melanogaster* flies at different densities and measured the effect this has on their post-infection survival and resistance to starvation. We find that flies housed at higher densities show greater mortality after being infected with bacterial pathogens, while also exhibiting increased resistance to starvation. Our results are more in line with the density-stress hypothesis that postulates a compromised immune system when hosts are subjected to high densities.

## Introduction

Maintaining a functional immune system, constitutively ready to tackle attacking pathogens imposes costs upon the host (McKean et al. 2008, Shudo and Iwasa 2001). Mounting an immune response after being challenged by a pathogen/parasite also requires considerable resource investment on part of the hosts leading to costs in terms of life-history and reproductive functions (Sheldon and Verhulst 1996, Lochmiller and Deerenberg 2000, Schmid-Hempel 2003). Host organisms are thus expected to modulate their immune function depending upon numerous factors, including pathogen prevalence, physiological state, access to compensatory resources, etc. (Lazzaro and Little 2009).

Risk of infection increases with increasing host population density because of higher between individual contact (Anderson and May 1981, Dwyer 1991). Under such conditions where there is a high risk of infection, hosts may prophylactically increase their level of immune function as a precaution towards impending infection (density-dependent prophylaxis hypothesis; Wilson and Reeson 1998). This prophylactic immune response can protect hosts from getting infected (viz. the protection against fungal and parasitoid infection in insects provided by melanized cuticle that develops in response to increased population density) or help hosts survive better after being infected with a pathogen.

High population density, in addition to increasing pathogen prevalence and infection risk, can also reduce per capita availability of resources and causes physiological stress. Since maintaining and deploying the immune system are both resource intensive process, reduced availability of resources and stressed physiological state can compromise functionality of the immune system (Wakelin 1989, Lazzaro and Little 2009). High population density can therefore make hosts immuno-compromised and increase their susceptibility to invading pathogens and parasites (crowding stress hypothesis; Steinhaus 1958). The exact observed effect of manipulating host population density on the host immune phenotype will therefore probably depend upon the exact mechanistic interaction occurring in the host-pathogen system under investigation, and also on the severity of the density manipulation implemented in the study. For example, since pathogens are themselves dependent upon host resources to fuel their systemic growth and virulence, limiting host access to resources can have a complex, pathogen-dependent effect on the outcome of infection (Pike et al. 2019).

Insects, and other invertebrates, have been actively used as models to study the effect of host population density on host immune function (Wilson and Cotter 2009). Many insect populations in the wild go through major cycles of population fluctuations making insects a relevant study system for such questions. Habitat destruction and climate change induced changes in insect population densities make the matter more relevant not just to scientists with entomological interests but also to public in general (Wagner et al. 2021). Increasing density has been found to increase immune function and pathogen resistance in *Mythimma separata* (Mitsui and Kunimi 1988, Kunimi and Yamada 1990), *Spodoptera exempta* (Resson et al. 1998, Resson et al. 2000, Wilson et al. 2001), *Spodoptera littoralis* (Cotter at al. 2004), *Mamestra brassicae* (Goulson and Cory 1995), *Tenebrio molitor* (Barnes and Siva-Jothy 2000), *Schistocerca gregaria* (Wilson et al. 2002), *Bombus terrestris* (Ruiz-Gonzalez et al. 2009), *Anabrus simplex* (Bailey et al. 2008)) and a few other insect species. These studies include experiments on a wide range of insect taxa, using different types of pathogens (virus, bacteria, and fungi), and manipulating density at various stages of host life history (larva, nymphs, and adults). A few studies although have failed to observe any density-dependent increase in immune function in certain insects, for example in *Zootermopis angusticollis* (Pie et al. 2005), *Gryllus texensis* (Adamo 2006), *Lymantria dispar* (Reilley and Hajek 2008), and *Danaus plexippus* (Lindsey et al. 2009).

There is limited knowledge on how population density, crowding, or social environment affects immune phenotypes in adult *Drosophila melanogaster*. Leech and colleagues (2019) compared flies housed individually versus flies housed in single sex pairs and demonstrated that being housed in pairs can improve post-infection survival in an age and pathogen specific manner, although this result is best interpreted in light of presence/absence of social contact (Bailey and Moore 2018) rather than in the paradigm of density manipulation. Kapila and colleagues (2021) have shown that adult flies that have survived crowded larval conditions tend to perform better when infected, but only for certain pathogens.

In the present study, using a lab adapted, outbred population of *Drosophila melanogaster*, we tested for the effect of manipulating adult density on adult immune function. We subjected adult flies to different densities (see METHODS for more details) and measured their immune function in form of post-infection survival after being infected with bacterial pathogens. Previous studies have demonstrated that housing *Drosophila melanogaster* adults in different group sizes can produce varied physiological responses. For example, under laboratory conditions, *Drosophila melanogaster* adults housed at different group sizes exhibit correlated changes in life history (Nandy et al. 2016; Leech et al. 2017) and sexually selected (Bretman et al. 2010, 2011; Nandy and Prasad 2011) traits, with effects extending into the next generation (Dasgupta et al. 2016, 2019). Flies subjected to crowding early in adult life have reduced lifespan (Graves and Meuller 1993; Joshi et al. 1998) and altered age-independent mortality rate (Joshi and Meuller 1998). Manipulating adult density also influence flies’ dispersal capabilities (Mishra et al. 2018). These results suggest that *Drosophila melanogaster* is a suitable model system to test for effects of manipulating adult density on adult immune function.

In addition to measuring the effect of different densities on immune function, we also measured resistance to starvation (i.e., longevity without access to food but allowed access to water) of the host after being subjected to different densities. Although simple and intuitive expectation suggest that resistance to starvation will be compromised following exposure to stressful environments, it has been suggested that exposure to high density can potentially prime organisms towards being more stress resistant and increase their survival under starving conditions (Rion and Kawecki 2007). Our results indicate that organisms conditioned at higher densities exhibit greater post-infection mortality depending upon the identity of the pathogen and the severity of density manipulation, but independent of host sex. Resistance to starvation is increased at the highest density, but only in female flies.

## Materials and Methods

### Study system and general fly handling

Flies from <u>B</u>lue <u>R</u>idge <u>B</u>aseline 2 (BRB2) population were used in experiments reported here. Originally established by hybridizing 19 iso-female lineages (themselves established from wild-caught females) (Singh et al. 2015), BRB2 has been maintained as a large (census size of approximately 2800 adults), outbred population for about 200 generations by the time these experiments were carried out. Every generation, eggs are collected from the population cage and distributed into glass vials (25 mm diameter × 90 mm height) with 6-8 mL of banana-jaggery-barley-yeast food medium at a density of 60-80 eggs per vial; 40 such vials are set up in total. The vials are incubated at 25 °C, 50-60% RH, with a 12:12 LD cycle. Eggs develop into adults in 9-10 days, and 12 days after egg collection (day of egg collection counted as day 1) all adults are transferred to a plexiglass cage (25 cm length × 20 cm width × 15 cm height) and provided with fresh food medium supplemented with *ad libitum* live yest paste for two days. On day 14, fresh food is provided to the cage and 18 hours later eggs are collected off the same to initiate the next generation. BRB2 is thus maintained on a 14-day discrete generation cycle.

### Deriving flies for experiments

Experimental flies were reared and handled under conditions identical to the regular maintenance of the BRB2 population. Eggs were collected at a density of 60-80 eggs per vial, with each vial having 8 mL of standard food medium; 80 such vials were set up. On day 12 post-egg collection, adult flies from all vials were pooled together and randomly sorted into different density treatments (see below) under light CO_2_ anesthesia. Adults in different density treatments were housed in glass vials with 1.5-2 mL of standard food medium (same amount of food was provided to flies of all treatments) till the day of immune/starvation assay (day 14 post-egg collection).

### Experimental design

#### Experiment 1: Effect of density, 8 adults vs. 32 adults, on immune function and starvation resistance

On day 12 post-egg collection, adult flies were sorted into fresh food vials (with 1.5-2 mL of food medium) at densities of 8 individuals or 32 individuals in each vial, in 1:1 sex ratio. The flies were held in these vials for two days, the *conditioning* period. After the conditioning the flies were subjected to infections (or, starvation) as described below, and housed at density of 4 males and 4 females per vial. For immunity assays, 20 infection vials were set up per density treatment and 10 sham-infection vials were set up per treatment. The experiment was replicated twice with each pathogen. For starvation resistance assays, 20 vials were set up per density treatment, and the assay was replicated twice.

#### Experiment 2: Effect of density, 50 adults vs. 200 adults, on immune function and starvation resistance

On day 12 post-egg collection, adult flies were sorted into fresh food vials (with 1.5-2 mL of food medium) at densities of 50 individuals or 200 individuals in each vial, in 1:1 sex ratio. The flies were conditioned at these densities for two days. After the conditioning the flies were subjected to infections (or, starvation) as described below, and housed at density of 4 males and 4 females per vial. For immunity assays, 20 infection vials were set up per density treatment and 10 sham-infection vials were set up per treatment. The experiment was replicated twice with each pathogen. For starvation resistance assays, 20 vials were set up per density treatment, and the assay was replicated twice.

### Immune function assay

We measured post-infection survival of flies as a proxy for immune function. Flies were infected with two bacterial pathogens separately: *Erwinia c. carotovora* and *Enterococcus faecalis*. 5 mL lysogeny broth (Luria Bertani Broth, Miller, HiMedia) was inoculated with a stab of bacterial glycerol stock and incubated overnight at 37 °C with aeration. Secondary culture was established by inoculating 10 mL lysogeny broth using 100 uL of this culture and allowed to grow till the culture was confluent. The bacterial cells were then pelleted down via centrifugation and re-suspended in sterile MgSO_4_ buffer at 1.0 OD_600_. Flies were infected by pricking them in the thorax under light CO_2_ anesthesia with 0.01 mm Minutein pins (Fine Scientific Tools, USA) dipped in the bacterial slurry. Sham-infections were done similarly except that the pins were dipped into sterile MgSO_4_ buffer. Flies were then placed in fresh food vials. They were shifted to fresh food vials 72 hours post-infection. Vials were monitored every 4-6 hours and mortalities were recorded for 120 hours post-infection. For all infection experiments, sample size per density treatment was 80 males and 80 females for infections and 40 males and 40 females for sham-infections.

### Starvation resistance assay

For assaying starvation resistance, flies were placed into glass vials with 2 mL of non-nutritive agar (2%) gel. Flies were transferred to new vials every 2-3 days, and checked for mortality every 8-10 hours till every fly was dead. For all starvation resistance experiments, sample size per density treatment was 80 males and 80 females.

### Statistical analysis

Survival analysis was done in R (R Core Team 2019, v3.6.2) using the packages *survival* (Therneau 2015, v2.38) and *coxme* (Therneau 2020, v2.2-16). Survivorship data from infection experiments were analyzed using a mixed-effects Cox Proportional Hazards model, with survival modeled as:

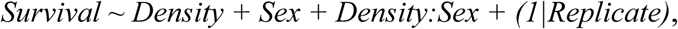

where ‘Density’, ‘Sex’, and their interaction are modeled as fixed factors and ‘Replicate’ is modeled as a random factor. Since very little and negligible mortality was observed in the sham infection controls, only data from infected flies was used in this analysis. Survivorship data from starvation resistance experiments were analyzed similarly using a mixed-effects Cox Proportional Hazards model, with survival modeled as:

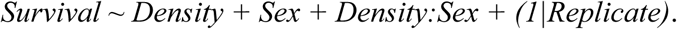

Survival curves were plotted using the package *survminer* (Kassambara et al. 2019, 0.4.6).

## Results

In the first experiment, we quantified the effect of housing flies in groups of 8 vs. 32 flies (1:1 sex ratio) per vial for 48 hours in terms of immune function and resistance to starvation; post-infection survival when infected by *Enterococcus faecalis*, and *Erwinia c. carotovora* was used as proxy of immune function (figure 1 and table 1). No mortality was recorded in either density treatment during the 48-hour conditioning period. Flies housed at the density of 32 flies per vial had reduced survival compared to flies housed at a density of 8 flies per vial when infected with *E. c. carotovora* (hazards ratio, 95% CI: 1.359, 1.012-1.827). Housing density had no significant effect on survival post infection with *E. faecalis* (hazards ratio, 95% CI: 1.257, 0.941-1.679). Flies from both density treatments had similar survival when subjected to starved conditions (hazards ratio, 95% CI: 1.193, 0.956-1.489). Sex of the flies was a significant determinant of survival in case of flies infected with *E. c. carotovora*; males were more susceptible to infection compared to females (hazards ratio, 95% CI: 1.977, 1.481-2.638). Host sex did not affect mortality in case of flies infected with *E. faecalis* (hazards ratio, 95% CI: 1.013, 0.751-1.481). Males were also more susceptible to starvation compared to females (hazards ratio, 95% CI: 1.868, 1.489-2.344). Any effect of interaction between density and sex was not observed in either measures of immune function or in resistance to starvation.

**Figure 1.**
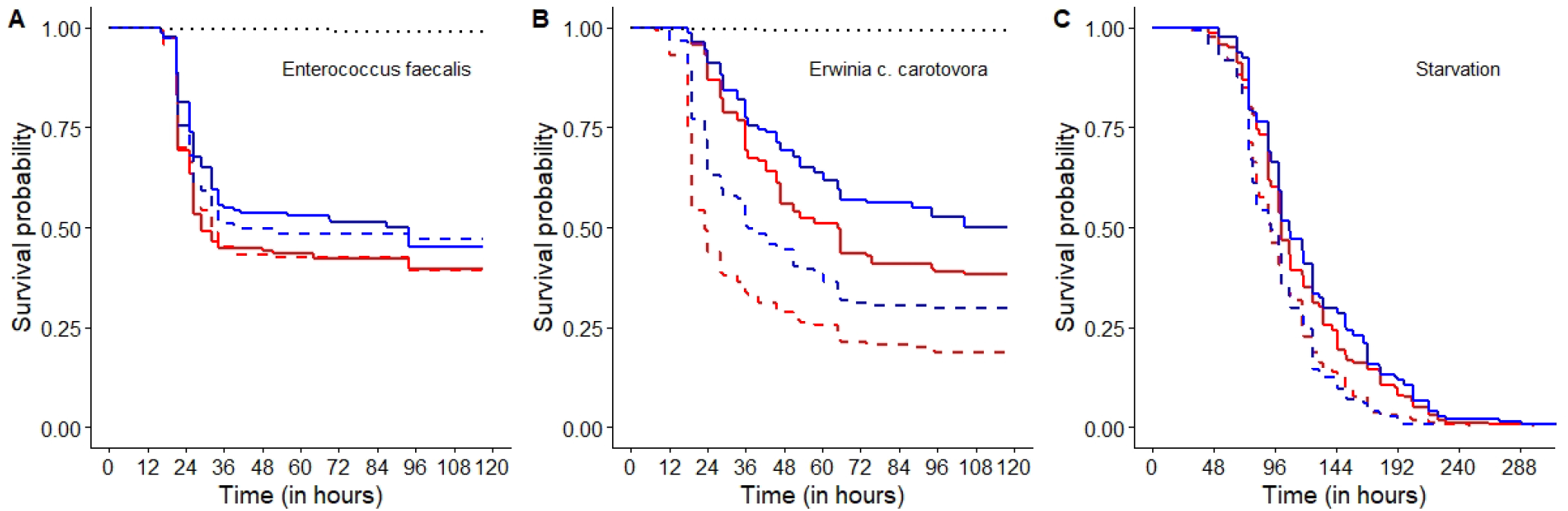
Survival curves for flies conditioned for 48 hours at densities of 8 vs. 32 flies per vial when (A) infected with *Enterococcus faecalis*, (B) infected with *Erwinia c. carotovora*, and (c) under starved conditions. Flies conditioned in groups of 32 are represented in ‘red’ and flies conditioned in groups of 8 are represented in ‘blue’; ‘solid’ lines represent female flies and ‘dashed’ lines represent male flies. The ‘black, dotted’ lines in (A) and (B) represent the flies subjected to sham-infections, all treatments pooled.

**Table 1.**
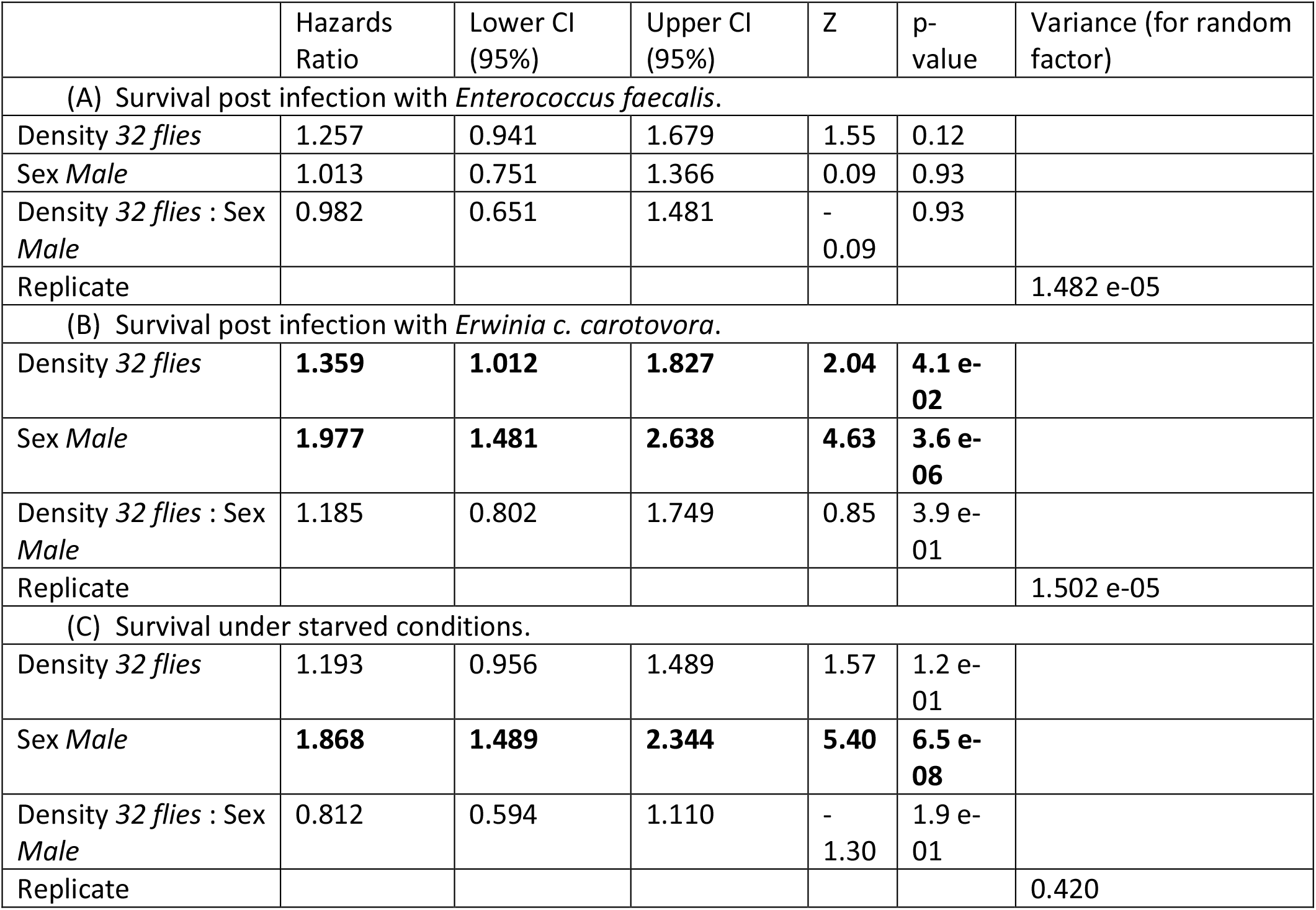
Output of mixed-effects Cox proportional hazards analysis of flies conditioned for 48 hours at densities of 8 vs. 32 flies per vial when (A) infected with *Enterococcus faecalis*, (B) infected with *Erwinia c. carotovora*, and (C) under starved conditions. Hazard ratios are relative to the default level for each factor, which is set at 1. The default level for “Density” is ‘8 flies’, and the default level for “Sex” is ‘Female’. Hazard ratio greater than 1 implies reduced survival compared to the default level.

|  | Hazards Ratio | Lower CI (95%) | Upper CI (95%) | Z | p-value | Variance (for random factor) |
| --- | --- | --- | --- | --- | --- | --- |
| (A) Survival post infection with *Enterococcus faecalis*. |  |  |  |  |  |  |
| Density 32 flies | 1.257 | 0.941 | 1.679 | 1.55 | 0.12 |  |
| Sex Male | 1.013 | 0.751 | 1.366 | 0.09 | 0.93 |  |
| Density 32 flies : Sex Male | 0.982 | 0.651 | 1.481 | -0.09 | 0.93 |  |
| Replicate |  |  |  |  |  | 1.482 e-05 |
| (B) Survival post infection with *Erwinia c. carotovora*. |  |  |  |  |  |  |
| Density 32 flies | **1.359** | **1.012** | **1.827** | **2.04** | **4.1 e-02** |  |
| Sex Male | **1.977** | **1.481** | **2.638** | **4.63** | **3.6 e-06** |  |
| Density 32 flies : Sex Male | 1.185 | 0.802 | 1.749 | 0.85 | 3.9 e-01 |  |
| Replicate |  |  |  |  |  | 1.502 e-05 |
| (C) Survival under starved conditions. |  |  |  |  |  |  |
| Density 32 flies | 1.193 | 0.956 | 1.489 | 1.57 | 1.2 e-01 |  |
| Sex Male | **1.868** | **1.489** | **2.344** | **5.40** | **6.5 e-08** |  |
| Density 32 flies : Sex Male | 0.812 | 0.594 | 1.110 | -1.30 | 1.9 e-01 |  |
| Replicate |  |  |  |  |  | 0.420 |

In the second experiment, we quantified the effect of housing flies in groups of 50 vs. 200 flies (1:1 sex ratio) per vial for 48 hours in terms of immune function and resistance to starvation (figure 2 and table 2) after being conditioned at different densities. The aim was to rule out the possibility that the reason no effect of housing density was observed in experiment 1, except in context of post-infection survival when infected with *E. c. carotovora*, was that the density treatments were not sufficiently different from one another in terms of severity. Although no mortality was observed during the 48-hour conditioning period in flies housed at density of 50 per vial, flies housed at density of 200 per vial exhibited mortality during conditioning; about 5-8% of flies of both sexes died in each vial. Flies housed at a density of 200 flies per vial died significantly faster compared to flies housed at a density of 50 flies per vial when infected with both *E. faecalis* (hazards ratio, 95% CI: 1.313, 1.002-1.721) and *E. c. carotovora* (hazards ratio, 95% CI: 1.472, 1.061-2.043). Males died faster when infected with *E. c. carotovora* (hazards ratio, 95% CI: 2.120, 1.538-2.992); sex had no effect on survival post infection with *E. faecalis* (hazards ratio, 95% CI: 0.928, 0.699-1.230). Although there was an overall effect of density on resistance to starvation, where flies housed at the high density died slower compared to flies housed at lower density (hazards ratio, 95% CI: 0.770, 0.616-0.963), survival under starved condition was predominantly affected by interaction between density treatment and sex, with the difference between densities being more prominent in case of female flies (figure 2(C)). Overall males died faster when subjected to starvation (hazards ratio, 95% CI: 1.681, 1.344-2.103).

**Figure 2.**
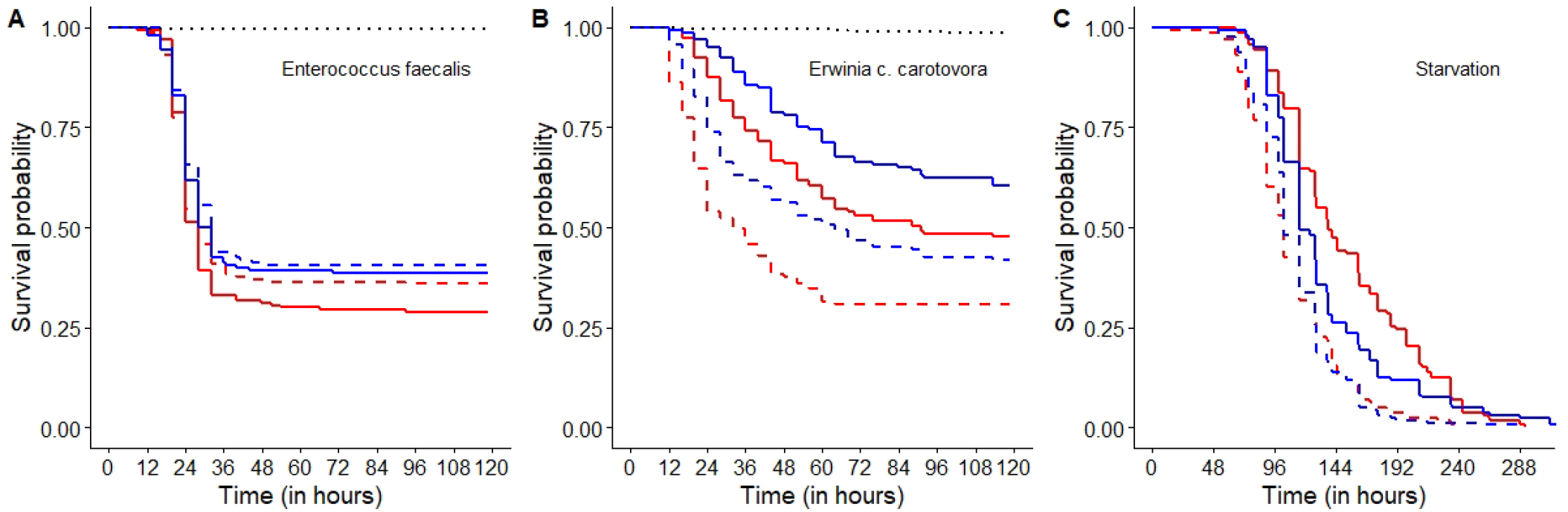
Survival curves for flies conditioned for 48 hours at densities of 50 vs. 200 flies per vial when (A) infected with *Enterococcus faecalis*, (B) infected with *Erwinia c. carotovora*, and (c) under starved conditions. Flies conditioned in groups of 200 are represented in ‘red’ and flies conditioned in groups of 50 are represented in ‘blue’; ‘solid’ lines represent female flies and ‘dashed’ lines represent male flies. The ‘black, dotted’ lines in (A) and (B) represent the flies subjected to sham-infections, all treatments pooled.

**Table 2.** Output of mixed-effects Cox proportional hazards analysis of flies conditioned for 48 hours at densities of 8 vs. 32 flies per vial when (A) infected with *Enterococcus faecalis*, (B) infected with *Erwinia c. carotovora*, and (C) under starved conditions. Hazard ratios are relative to the default level for each factor, which is set at 1. The default level for “Density” is ‘50 flies’, and the default level for “Sex” is ‘Female’. Hazard ratio greater than 1 implies reduced survival compared to the default level.

|  | Hazards Ratio | Lower CI (95%) | Upper CI (95%) | Z | p-value | Variance (for random factor) |
| --- | --- | --- | --- | --- | --- | --- |
| (A) Survival post infection with *Enterococcus faecalis*. |  |  |  |  |  |  |
| Density 200 flies | **1.313** | **1.002** | **1.721** | **1.98** | **0.048** |  |
| Sex Male | 0.928 | 0.699 | 1.230 | - 0.52 | 0.600 |  |
| Density 200 flies : Sex Male | 0.926 | 0.628 | 1.366 | - 0.39 | 0.700 |  |
| Replicate |  |  |  |  |  | 0.011 |
| (B) Survival post infection with *Erwinia c. carotovora*. |  |  |  |  |  |  |
| Density 200 flies | **1.472** | **1.061** | **2.043** | **2.32** | **2.1 e-02** |  |
| Sex Male | **2.120** | **1.538** | **2.922** | **4.59** | **4.5 e-06** |  |
| Density 200 flies : Sex Male | 1.089 | 0.709 | 1.672 | 0.39 | 6.9 e-01 |  |
| Replicate |  |  |  |  |  | 0.526 |
| (C) Survival under starved conditions. |  |  |  |  |  |  |
| Density 200 flies | **0.770** | **0.616** | **0.963** | - **2.29** | **2.2 e-02** |  |
| Sex Male | **1.681** | **1.344** | **2.103** | **4.56** | **5.2 e-06** |  |
| Density 200 flies : Sex Male | **1.377** | **1.008** | **1.881** | **2.01** | **4.4 e-02** |  |
| Replicate |  |  |  |  |  | 0.023 |

## Discussion

In this study we tested for the effect of increasing adult density on immune function and starvation resistance of *Drosophila melanogaster*. We find that increasing density makes flies more susceptible to infection with pathogenic bacteria. Increasing density, in the other hand, helps prolong lifespan of flies under starved conditions. Many studies in the past have tested for the effect of host population density on host immune function. The density-dependent prophylaxis (DDP) hypothesis suggests, that due to anticipated increase in risk of infection when population densities increase, individual organisms upregulate their immune system as a prophylactic counter measure (Wilson and Reeson 1998). Alternatively, the crowding-stress hypothesis proposes that at high population density organism’s physiological capabilities are compromised due to stress and resource limitation leading to individuals having sub-optimal immune function (Steinhauss 1958). Our results, therefore, seem to align more with the crowding-stress hypothesis.

Results from experiments aiming to characterize the effect of population density on immune function have been variable. One possible reason for this variance is that the different studies use different measures of immune function. Effect of phenotypic and environmental manipulations often have dissimilar and opposing effects on different sub-organismal readouts of immune function (Adamo 2004a, Adamo 2004b). For example, in *Spodoptera littoralis*, melanic larvae reared in high densities exhibit increased hemolymph phenoloxidase activity compared to non-melanic and solitary larvae, but have reduced hemolymph antimicrobial activity levels (Cotter et al. 2004). Whereas in *Schistocerca gregaria*, individuals reared under crowded conditions have increased antimicrobial activity compared to individuals raised in solitary, without any change in phenoloxidase activity or cellular encapsulation response (Wilson et al. 2002). Additionally, if different immune defense mechanisms are affected differently by density manipulation, the observed effect of density on outcome of infection can vary depending upon the pathogen used to challenge the hosts. As a potential solution we, therefore, used post-infection survival against two different entomopathogenic bacteria as proxy of immune function in our experiments.

In the experiments reported in this paper, we subjected adult flies to two different density ranges, and then quantified their resistance against bacterial pathogens and starvation. The idea behind using two different density ranges was to test for the effect of increasing density with and without changing the amount of physiological stress suffered by the focal organisms (Goulson and Corey 1995). In the experiments comparing between 8 adults vs. 32 adults per vial (experiment 1), there was no differential mortality during the conditioning period, or even after that (shams died equally and negligibly in both treatments; figure 1(A) & 1(B)). But in experiments comparing between 50 adults vs. 200 adults per vial (experiment 2), mortality was observed during the two-day conditioning period in case of adults held at 200 flies per vial but not in adults held at 50 flies per vial; post-conditioning mortality was again not different between density treatments (shams died equally and negligibly in both treatments; figure 2(A) & 2(B)).

To make our immune function experiments generalizable, we tested survival of the focal flies when infected with two bacteria pathogens, one Gram negative (*Erwinia c. carotovora*) and one Gram positive (*Enterococcus faecalis*). Previous research into *Drosophila* immunity has established that the mechanisms employed to counter these two types of pathogens are significantly different, with some level of cross-talk (Buchon et al. 2014).

The results from our immunity experiments indicate an absence of DDP in response to adult crowding. When infected with *E. c. carotovora*, across both density range comparisons, flies housed at lower density had better post-infection survival compared to flies housed at higher density (figures 1(B) and 2(B)). For flies infected with *E. faecalis*, no effect of conditioning at different density was apparent in experiment 1 (comparison between conditioning at density 8 vs. 32 flies per vial; figure 1(A)), but in experiment 2, flies housed at the lower density (50 flies per vial) had better survival compared to flies housed at the higher density (200 flies per vial); figure 2(A). The effect of housing density on post-infection survival is expected to be due to the differences created during the conditioning window as during the experiments (post-infection) all adults were housed at equal density. Previous research has shown that change in density both before and after administration of pathogen can interact to influence immune function (Kumini and Yamada 1990). The canonical alternative to DDP hypothesis is the crowding stress hypothesis, which argues that higher density leads to physiological stress and lack of resources, and therefore should compromise immune function. The results from our immune function experiments with both pathogens seem to agree very well with this idea. The differences between results with E. c. carotovora and *E. faecalis* may be attributed to differential energetic costs of immunity against these two pathogens. Defense against *E. faecalis* may in fact be cheap: previous research has shown that when flies are reared on substandard diet, or prevented from feeding, their resistance against *E. faecalis* is affected much less compared to resistance against other pathogens (Ayres and Schneider 2009). Subjecting flies to larval crowding also has no effect on adults’ immune defense against *E. faecalis* (Kapila et al. 2021).

We did not observe any sex × density interaction in context of post-infection survival in case of both pathogens used here. Sexes differ in their life history and reproductive strategies, and are therefore expected to have different optimal investment towards immune function (Rolff 2002). One could therefore hypothesize that crowding stress can potentially affect the two sexes differently. But our results indicate that whatever effect density has on immune function is equivalent for both sexes. Interaction between sexes can influence immune function of insects (Rolff and Siva-Jothy 2002; Winterhalter and Fedorka 2009). In our experiments, we held the sexes together during the period of density conditioning, albeit at equal sex ratio. If inter-sexual interactions (such as mating rate and harassment) change in a density-dependent fashion, results from our immunity assay might be explained by such changes. In *Drosophila melanogaster* mating rate has a strong bearing on male immune function while female immune function is mostly determined by access to resources (McKean and Nunney 2005). Therefore, even when the observed phenotypic effect of increasing density is similar for both sexes, the effect can be, for each sex, driven by separate factors (mating rate in males and access to resources in females) that co-vary independently with changing density.

Increasing population density can have complex effects on an insect’s response to stress. Reduced access to resources should directly translate to compromised physiology and low systemic resource reserves leading to increased vulnerability to stress such as starvation. Exposure of adults to high density has been shown to compromise immediate mortality rate even when flies are returned to normal conditions (Joshi and Meuller 1998). Alternatively, exposure to increased density (and thereby shortage of resources) can prime flies hormetically to be better resistant to starvation (Rion and Kawecki 2007). Additionally, since sexes are known to be differentially susceptible to starvation stress (Chippindale et al. 1996; Harshman and Schmid 1998), the effect of increasing population density on starvation resistance may be sex specific. In our experiment comparing flies conditioned at densities of 8 versus 32 adults, we find no effect of density of starvation resistance, although we see a major effect of sex, confirming earlier results. In the follow-up experiment comparing flies housed at densities of 50 versus 200 adults, we see no effect of density on starvation resistance in males; but among females, flies that were housed at higher density exhibit a significantly increased resistance to starvation. Our observations are in line with the prediction that crowding stress can prime flies to be less susceptible to starvation (Rion and Kawecki 2007). Since female fitness is influenced by access to resources to a greater extent compared to males (Bateman 1948), it is not surprising that we observe high density induced priming against starvation only in case of female flies. Drosophila melanogaster females are known to suppress egg laying in response to increased adult density in culture vials (Pearl 1932, Barker 1973). This reduced investment towards reproduction can be an additional explanation for increased starvation resistance of females conditioned at higher density (200 adults per vial); flies adapted to starvation resistance often have reduced fecundity (Service et al. 1988).

To summarize, in our experiments we demonstrate an overall negative effect of increasing adult density on the flies’ ability to survive infection with bacterial pathogens, indicating that increased density compromises immune function; this observation agrees with the prediction from the crowding stress hypothesis (Steinhaus 1958). We also demonstrate that under certain conditions, increasing adult density can lead to a sex-specific increase in resistance to starvation, agreeing with the prediction of high density induced priming against future stressors (Rion and Kawecki 2007). Put together, these results indicate that increase in population density, and associated lack of resources, can subject insects to physiological stress resulting in major fitness differences; a result with far reaching consequences. It would help our understanding further if future studies aim to parse out the two different, yet simultaneously acting, features of density stress: crowding (having more conspecifics present around) from the effect of reduced resource availability.

## Acknowledgements

We thank Dr. Elio Sucena and Tania F T Paulo (Instituto Gulbenkian Ciência, Oeiras, Portugal) for lending us the *Erwinia c. carotovora* isolate, and Prof. Brian Lazzaro (Dept. of Entomology, Cornell University, Ithaca, USA) for lending us the *Enterococcus faecalis* isolate used in this study. We also thank Aparajita Singh and Rochishnu Dutta for their constructive comments on earlier drafts of the manuscript.

